# *Pristionchus* nematodes occur frequently in diverse rotting vegetal substrates and are not exclusively necromenic, while *Panagrellus redivivoides* is found specifically in rotting fruits

**DOI:** 10.1101/324996

**Authors:** Marie-Anne Félix, Michael Ailion, Jung-Chen Hsu, Aurélien Richaud, John Wang

## Abstract

The lifestyle and feeding habits of nematodes are highly diverse. Several species of *Pristionchus* (Nematoda: Diplogastridae), including *Pristionchus pacificus*, have been reported to be necromenic, i.e. to associate with beetles in their dauer diapause stage and wait until the death of their host to resume development and feed on microbes in the decomposing beetle corpse. We review the literature and suggest that the association of *Pristionchus* to beetles may be phoretic and not necessarily necromenic. The view that *Pristionchus* nematodes have a necromenic lifestyle is based on studies that have sought *Pristionchus* only by sampling live beetles. By surveying for nematode genera in different types of rotting vegetal matter, we found *Pristionchus* spp. at a similar high frequency as *Caenorhabditis*, often in large numbers and in feeding stages. Thus, these *Pristionchus* species may feed in decomposing vegetal matter. In addition, we report that one species of *Panagrellus* (Nematoda: Panagrolaimidae), *Panagrellus redivivoides*, is found in rotting fruits but not in rotting stems, with a likely association with *Drosophila* fruitflies. Based on our sampling and the observed distribution of feeding and dauer stages, we propose a life cycle for *Pristionchus* nematodes and *Panagrellus redivivoides* that is similar to that of *C. elegans*, whereby they feed on the microbial blooms on decomposing vegetal matter and are transported between food patches by coleopterans for *Pristionchus* spp., fruitflies for *Panagrellus redivivoides* and isopods and terrestrial molluscs for *C. elegans*.

## Introduction

The lifestyle and feeding habits of members of the nematode phylum are highly diverse. A single species may express diverse life cycles through production of free-living and host-associated stages. Many species of the nematode genus *Pristionchus* (Nematoda: Diplogastridae), including the most studied *Pristionchus pacificus*, have been recently considered to be associated with beetles in a necromenic lifestyle, i.e. to associate with beetles in the nematode dauer diapause stage and wait until the death of the beetle to resume development and feed on microbes in the decomposing beetle corpse [1]. The necromenic lifestyle of *Pristionchus* nematodes now appears as a qualifier for the species *P. pacificus* [2,3]. Thus, *Pristionchus* have been considered to share some attributes of parasitic nematodes and to have a very different lifestyle from the free-living nematode *C. elegans* [4].

Yet, to our knowledge, carcasses of naturally dead beetles have not been looked for and examined to assess whether *Pristionchus* feeds on them. Recent field studies have looked for *Pristionchus* in live beetles, and to a lesser extent in soil [5–8]. We detail below in three points what has been documented about the association of *Pristionchus* with beetles.

- 1) Many *Pristionchus* species including *P. pacificus* can be found on beetles [6,8–14]. (Note that a subclade of *Pristionchus* species has recently been found in fresh tropical figs in association with their pollinating wasps [15], in the same way that some *Caenorhabditis* species are found in other specific substrates [16] - we will not consider this fig subclade here.)

- 2) *Pristionchus* can be found on live beetles in the dauer stage [12], based on collecting 114 live *Geotrupes stercorosus* beetles in a forest near Tübingen, Germany. The authors found a total of 17 *Pristionchus* individuals, 71 *Koerneria* (a genus close to *Pristionchus*), 466 other diplogastrids, 3927 *Pelodera* (Rhabditidae) and 508 parasitic Spirurida. *Pelodera* dauer larvae were present in 56 out of 114 *Geotrupes* beetles, with up to hundreds of *Pelodera* dauer larvae per beetle. *Pristionchus* was comparatively rarer: eight of 114 beetles yielded one *Pristionchus* individual, one carried two individuals and one had seven individuals, all in the dauer stage. *P. pacificus* is rare in Europe [9]. These beetle-associated *Pristionchus* were *P*. sp. 6, *entomophagus*, and *lheritieri*. In addition, a survey of 4242 beetles from Germany, France, Spain, Switzerland again suggested that related *Pristionchus* spp. are found in the dauer stage on beetles [9].

- 3) Finally, the data supporting the conclusion that *Pristionchus* non-dauer stages feed on carcasses of beetles come from experiments where beetles were killed on a Petri dish containing *E. coli* in the laboratory (e.g., [9], and for *P. pacificus*: [6]). When a beetle is killed on a Petri dish, some of the bacterial species that are inside the gut or on the surface of the animal will proliferate using the beetle or the Petri dish as food and *Pristionchus* can then feed on them or *E. coli* (cf. Fig 1 in [8]). In these experiments, *Pristionchus* appeared only after 7-10 days, suggesting that they took time to exit the dauer stage [9]. The delay may be explained by the fact that some beetles may secrete chemicals that prevent their associated nematode species from exiting the dauer stage [17]. Because this delay is longer than the generation time in this environment (3-4 days), it is not possible to assess either the number of individuals or the stage at which they were on the beetle. Similarly, if the beetles are killed by the experimenter and then placed in a soil environment, *Pristionchus* will proliferate in the vicinity of the dead beetle only after 7 days [18]. These data suggest that *Pristionchus* may adopt a necromenic lifestyle by consuming the progeny of bacteria that were on the living host beetle, but they do not address whether this is an event occurring in the wild, and if so, how frequent it is for *Pristionchus*. The claim of a necromenic relationship does not take into account the possibility that the nematodes may disembark from their insect host well before it dies, should they encounter some food source. Necromeny may take place occasionally, but beetle corpses may be a very minor source of food for *Pristionchu*s.

**Fig 1.**
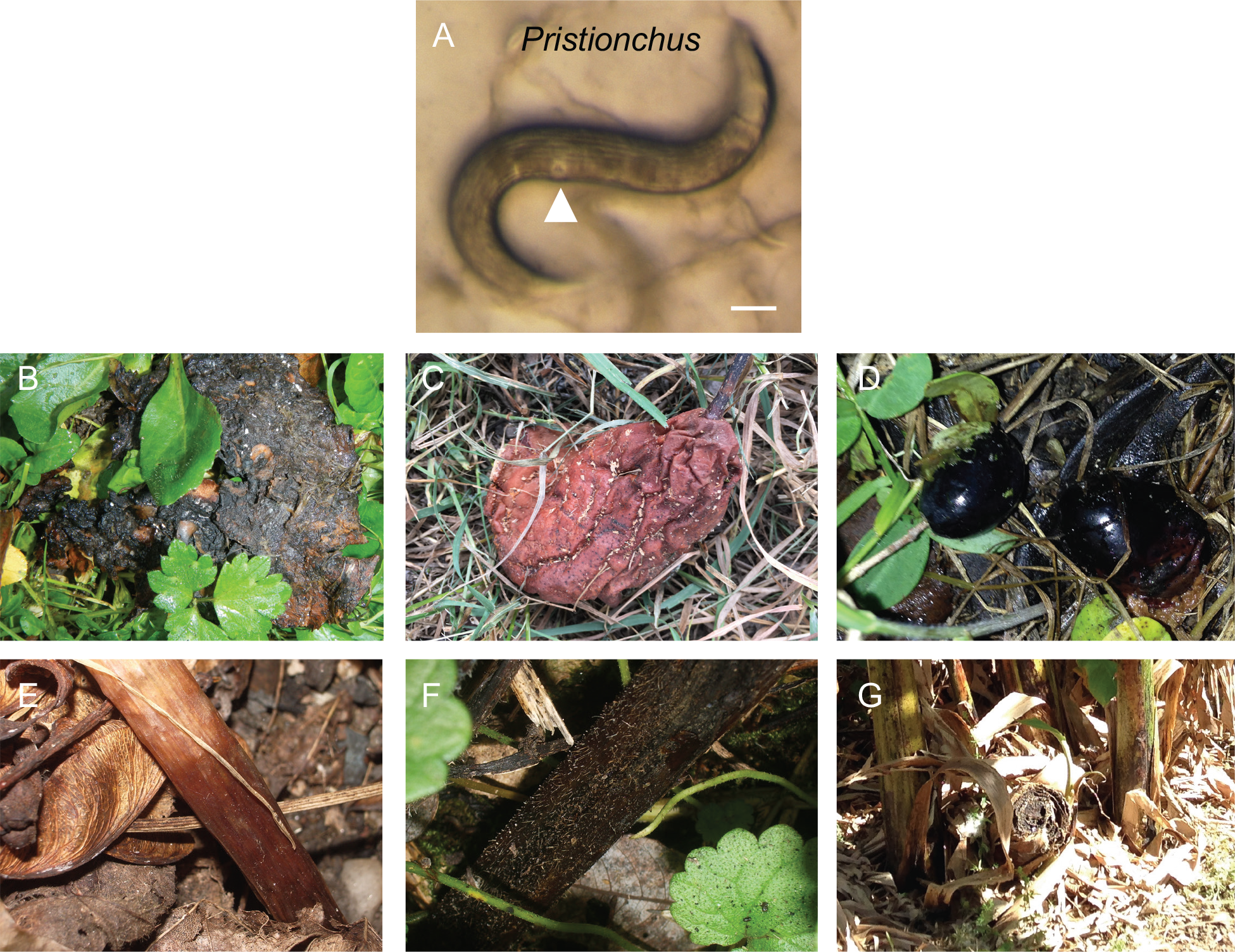
Rotting fruit and stem samples containing *Pristionchus*. (A) *Pristionchus* adult female or hermaphrodite from rotting apple O824 from Orsay. The cuticle with longitudinal ridges and the pore-shaped vulva (arrowhead) are characteristic of *Pristionchus*. Bar: 100 μm. (B) Apple O1194 in Orsay orchard, France with large (>1,000 individuals) populations including feeding individuals of *Pristionchus* sp. and *Caenorhabditis elegans*. (C) Pear CZ12 in Prague, Czech Republic with *Pristionchus* and *Panagrellus*. (D) Olives F8 near Firenze, Italy with *Pristionchus* sp. (E) *Arum* stem B09-6 in a wood near Le Blanc, France with *Pristionchus* and *Caenorhabditis*. (F) Stem S156 in a wood near Santeuil, France with a feeding population of several thousand *Pristionchus* individuals. (G) Banana pseudostem S9 in Palermo Botanical Garden, Italy, yielding *Pristionchus* and *C. elegans*.

By contrast, older literature on *Pristionchus* species had associated them with a number of substrates. Sudhaus and Fürst van Lieven [19] reviewed this systematic work on species in this genus and summarized the habitat as “mostly decaying plants, associated with insects”. Specifically, *Pristionchus* species were described from soil, humus, compost, moss, “diseased” stem, coffee berry, rotting bulbs of *Allium vineale*, damaged roots of coffee and garlic, around roots of several species, rotten potatoes, rotten wood, and decomposing fungi. Additionally, *Pristionchus* species were described to be associated with a wider range of insects than just beetles: termites (two species), *Ostrinia* (Lepidoptera), dead insects, and finally beetles (three species). The spectrum of habitats thus appears much wider than the recent literature indicates.

It is interesting to compare the recent history of research on *Pristionchus* ecology with what is known of the ecology of the model organism *Caenorhabditis elegans* and other *Caenorhabditis* species. The history of collecting *Caenorhabditis* has been different, as are the conclusions. *Caenorhabditis* nematodes have been found to be rare in soil, but abundant in rotting vegetal matter, such as fruits and soft plant stems. In addition, *C. elegans*, *C. briggsae* and *C. remanei* have been found on terrestrial molluscs, isopods and millipedes [20–27]. In laboratory experiments, the nematodes were shown to be capable of climbing on or off these invertebrates [20,21,27–30]. Perhaps anecdotally, *C. nouraguensis* were found on cockroaches in a tropical forest and *C. tropicalis* once on a beetle [16]. Some species, such as *C. japonica*, have an apparently specific carrier insect [31–34]. These associations are considered to be mostly phoretic or may correspond to occasional “facultative necromeny” [35], but this possibility has not been considered for *Pristionchus* spp.

We here provide data on nematode genera found in rotting vegetal matter, focusing on the genera *Pristionchus* and *Panagrellus*. By surveying different types of rotting vegetal and fungal substrates, we found diverse *Pristionchus* spp. at a similar frequency as *Caenorhabditis*, often in high numbers and in feeding stages in rotting fruits and stems. In addition, we report that a single species of *Panagrellus* (Nematoda: Panagrolaimidae), which we identify as *Panagrellus redivivoides*, is found in rotting fruits but not in rotting stems and appears to be associated with *Drosophila* fruitflies.

## Materials and Methods

### Sample handling and nematode genus identification

Collected samples were placed onto standard *C. elegans* Normal Growth Medium agar plates [36], previously seeded with *Escherichia coli* strain OP50 in the center of the plate. The samples were spread around the bacterial lawn. 1-2 ml water or M9 solution were added to humidify the samples [37].

For surveys in S1 Table, all plates were examined regularly under the dissecting microscope: in the most stringent surveys (Orsay, Santeuil), observations were made several times within the first hours, once or twice on the next two days, and at least on days 4 and 7. Nematodes that crawled out of the sample were identified to the genus or family level by morphological criteria, under the dissecting microscope with trans-illumination and sometimes further by Nomarski microscopy [37]. *Caenorhabditis*, *Pristionchus*, *Panagrellus*, *Oscheius*, *Mesorhabditis* species were identified as described in [26,37] (see pictures and drawings therein; and also [38,39] for identification keys). Our previous developmental evolution work provided us with experience on these various genera [40–48] and all strains then tested by rDNA sequencing confirmed correct identification to the genus level (S2 Table). In addition, *Caenorhabditis* were systematically identified to the species level as indicated in [26,37,49]. Given a generation time for these nematodes of about 3-5 days at 20°C, the number of individuals and developmental stages were noted for samples that were less than 48 hours from collection time.

These surveys were primarily aimed at studying *Caenorhabditis* populations, as reported in [22,25,26,49]. *Caenorhabditis* individuals tend to rapidly exit the sample to colonize the *E. coli* lawn, as do *Pristionchus*, *Panagrellus* and most other rhabditids. Some genera are more difficult to survey, either because they have a small body size, are present in small numbers, move slowly out of the sample or are less easy to recognize. The three genera *Caenorhabditis*, *Pristionchus* and *Panagrellus* that are highlighted in S1 Table, are those that may have been least missed in the samples. However, the number of samples that contained *Pristionchus* or *Panagrellus* spp. is likely an underestimate as they were less carefully looked for than *Caenorhabditis*.

Diplogastrids are easy to recognize by their short buccal cavity bearing strong teeth, the absence of a grinder in the basal bulb of the pharynx, the color pattern of the body stemming from the shape of female gonad arms generally bending back in diagonal towards the vulva, and the pore appearance of the vulva. *Pristionchus* are easy to recognize from other diplogastrids as adults, with their characteristic dumpyish body shape and cuticle with strong longitudinal ridges (Fig 1A). We also systematically checked that our morphological criteria corresponded to the *Pristionchus* genus by SSU rDNA sequencing of a subset of them (JU isolates, S2A Table).

*Panagrellus* species are recognized by their large body shape, light brown gut color (like *Caenorhabditis*), posterior vulva and viviparous reproduction (Fig 2A). Young first-stage larvae exit through the female vulva and gravid females may contain several dozens of embryos in their uterus. We also sequenced SSU rDNA for some of them (S2B Table).

**Fig 2.**
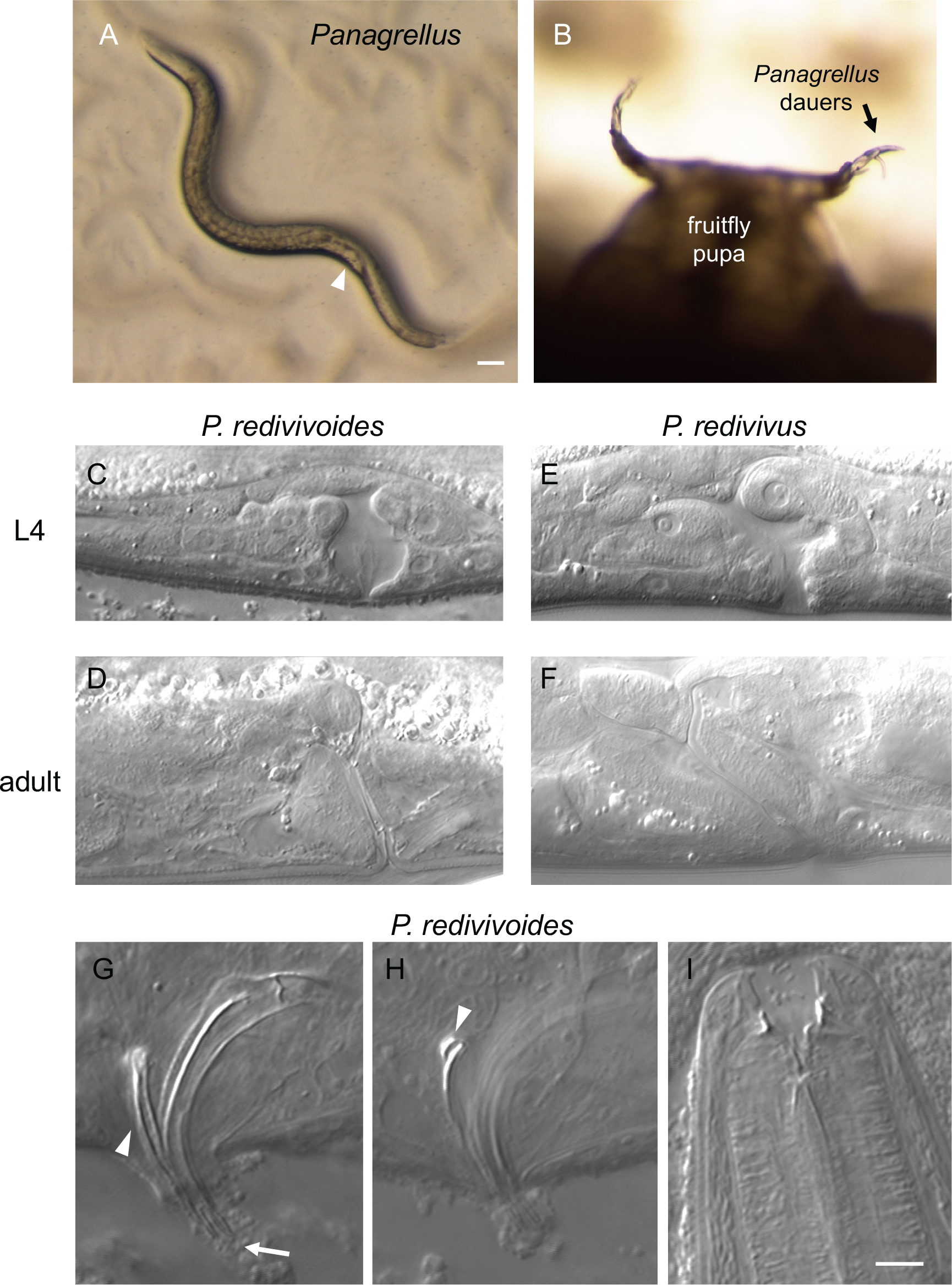
*Panagrellus* in rotting fruits. (A) *Panagrellus* adult female from rotting apple O801 from Orsay. The quite posterior vulva (arrowhead) and late-stage embryos accumulating in front of it are characteristic of *Panagrellus*. Bar: 100 μm. (B) *Panagrellus* dauers nictating on a *Drosophila* pupa in rotting pear B11-22 from Le Blanc. (C-F) Nomarski micrographs of L4 and adult vulvae of female *Panagrellus redivivoides* JU1476 (L4) and JU1798 (adult) and *Panagrellus redivivus* PS1163. Anterior is to the left, dorsal to the top for all panels, thus the uterus is on the left and the post-vulval sac is on the right in C-F. Most *Panagrellus* spp. display an anteriorly tilted vulva as shown for *P. redivivus*, while that of *P. redivivoides* is almost perpendicular to the ventral cuticle. (G) Spicule morphology of *P. redivivoides* (here strain JU1476). The forked ventral end of the spicules is indicated by an arrow, the gubernaculum by an arrowhead. (H) When in the proper focal plane, the dorsal side of the gubernaculum ends in the manner of a hook (arrowhead) (strain JU385). (I) Adult female mouth (here JU1055). Same scale for G-I. Bar: 5 μm.

Genera of other rhabditids can be *Pelodera*, *Auanema*, *Rhabditella*, *Pellioditis*, etc. In contrast to *Caenorhabditis* (or *Panagrellus*), their gut color is generally greyish/black, rather than brownish. Our notes are too scarce to distinguish them here and we placed them in a single category, with the exception of the ones for which we report 18S sequences.

*Oscheius* are very commonly found in rotting vegetal matter or soil and can be recognized by their greyish gut color, their lack of middle pharyngeal bulb, their thin female tail and their long and inflated rectum. The occurrence of *Oscheius* is likely underestimated as these animals are less striking, rarely reach large population sizes and develop more slowly than the above. They are here almost exclusively represented by small-size hermaphroditic *Oscheius* of the *Tipulae* and sometimes *Dolichura* subgroups (by contrast to large *Oscheius* of the *Insectivora* group; [45,50]).

*Panagrolaimus* are long and quite thin nematodes, with an only very slightly posterior vulva.

*Mesorhabditis* species are easier to find after one to two days on the plate and they may remain in the vicinity of the sample. They are recognized by their posterior vulva and dark body color. In our surveys, they were particularly looked for and isolated in 2015-17.

Animals of the *Protorhabditis/Prodontorhabditis/Diploscapter* clade [45,51] are small and tend to burrow in the agar and leave trails therein. Populations develop slowly and they can be missed.

*Rhabditophanes* has a black gut color and characteristically lay almost round eggs, instead of oval-shape ones. They were not systematically surveyed.

*Bunonema* individuals are easy to identify by their very small, spindle-shaped body, and a left-right asymmetric cuticle. They grow slowly and can easily be missed if their population is overwhelmed by other genera.

The “Other” category contains nematodes that are generally not bacterial eaters but fungi-eaters such as aphelenchs or parasitic nematodes that do not grow in our culture conditions.

### Culture and freezing

Strains were derived from single individuals in selfing species, or from a mated female or a male-female pair in male-female species, and cultured on standard *C. elegans* Normal Growth Medium agar plates. *Pristionchus* species were frozen with the *C. elegans* freezing protocol [36], sometimes adding 1 mM CaCl_2_ to the freezing solution. This protocol does not allow for a good retrieval of *Pristionchus* and *Panagrellus* nematodes. We recently adopted a DMSO-Dextran protocol that allows for better recovery (courtesy of Andre Pires da Silva), whereby a pellet of nematodes in M9 solution is resuspended in 7 ml of a mix of 1 g Dextran (Sigma D9260-500G), 1 ml DMSO and autoclaved H_2_O to 10 ml. The nematodes are incubated for 10 min at room temperature before being place in the −80°C freezer in styrofoam boxes.

### Mating tests

For the selfing *Pristionchus*, we crossed 5 hermaphrodites from the *P. pacificus* JU1102 strain [11] with 5 males of our new isolates and monitored the proportion of males in the progeny as an evidence of crossing. For two of them, BRC20259 and BR20261, we further checked whether these F1 males were fertile by crossing them to hermaphrodites of either parental strain. All crosses were positive. Two replicates of a control with JU1102 animals only did not yield a high percentage of males.

For *Panagrellus*, we crossed 3-5 L4 females with 3-5 males and monitored first and second generation progeny. A control with 5 L4 JU385 females (2 replicates) did not yield any progeny.

### PCR and sequence analysis of rDNA

In some surveys indicated in S2 Table, the nematodes were assigned to a genus using a molecular tag. The small subunit (SSU, 18S) of ribosomal DNA of *Pristionchus* isolates was amplified using primers SSU18A (5′-AAAGATTAAGCCATGCATG-3′) and SSU26R (CATTCTTGGCAAATGCTTTCG), and sequenced using SSU18A or SSU9R (AGCTGGAATTACCGCGGCTG), as in [5–9,52]; alternatively, as indicated in S2 Table, primers RHAB1350F (5′-TACAATGGAAGGCAGCAGGC) and RHAB1868R (5′-CCTCTGACTTTCGTTCTTGATTAA) were used.

The large subunit (LSU, 28S) of ribosomal DNA of *Panagrellus* isolates was amplified using primers D2A (ACAAGTACCGTGGGGAAAGTTG) and D3B (TCGGAAGGAACCAGCTACTA) as in [53].

The sequences were trimmed by visual inspection of the chromatograms, leaving the first four nucleotides of the downstream primer when present at the end of the sequence. A ‘N’ corresponds either to a low-quality sequence (including the possibility of a gap, especially when the same nucleotide occurred at consecutive positions) or a putative polymorphism of rDNA repeats. The sequences are available at Genbank with accession # (TO BE SUBMITTED).

The sequences were run through NCBI Blast (April-May 2018) with default parameters. In case of the apparent polymorphism in our *Panagrellus redivivoides* 28S sequences (AGTTGATCGGGTGTTGGCTTCGGY; cf. S2B Table, column P), the highest peak was used in the blast analysis. Differences with the closest sequence in the database were manually checked.

### Data analysis

The abundance index is defined on a Log10 scale as in [26]. An index of 1 corresponds to one to 10 individuals, 2 for 11 to 100, 3 for 10^2^ to 10^3^, 4 for 10^3^ to 10^4^, and 5 for > 10^4^.

Statistical analysis was performed in R version 3.4.1 [54].

## Results

### Sampling for *Caenorhabditis*

We sampled rotting vegetal matter, mostly fruits and stems from herbaceous plants (Fig 1), and occasionally flowers, leaves, and wood as well as a few other sample types, such as fungi, soil, humus mixes and occasionally larger invertebrates (S1 Table, S2 Table). Our primary goal was to isolate *Caenorhabditis* spp. Around 20 years ago, we first had sampled soil and found some *Pristionchus* (see S2A Table, first lines) but as we could not find *Caenorhabditis* we then focused on richer rotting vegetal matter [25]. In 2015-2017, we also sought to isolate *Mesorhabditis* spp. and thus included samples of humus and rotting leaves where species of this genus are easy to find. We present here all samples for which we systematically noted the different nematode genera. Some locations (in France) were sampled on many occasions while others have been sampled once.

*Caenorhabditis* was found in 34% of rotting fruit samples (n=556) and 41% of stem samples (n=204) (Fig 3, Table 1). As noted previously [26], we found them in these various samples either in a feeding stage or in the dauer diapause stage.

**Table 1.** Occurrence of *Caenorhabditis*, *Pristionchus* and *Panagrellus* nematodes in rotting fruits and stems from various locations around the world. Note that while *Caenorhabditis* was thoroughly searched for, the number of positive samples for *Pristionchus* and *Panagrellus* are underestimates, both because they may have been overlooked and because we may not have kept notes on batches of samples without *Caenorhabditis*. Detailed data for each sample are in S1 Table, with locations in France first in alphabetical order, then non-French locations.

| Location | Total # rotting fruits | # fruits with *Caenorh.* | # fruits with *Prist.* | # fruits with *Panag.* | Total # rotting stems | # stems with *Caenorh.* | # stems with *Prist.* | # stems with *Panag.* |
| --- | --- | --- | --- | --- | --- | --- | --- | --- |
| Crouy-sur-Ourcq, FR | 4 | 0 | 0 | 0 | 14 | 10 | 3 | 0 |
| Le Blanc & Indre | 62 | 18 | 9 | 24 | 17 | 2 | 7 | 0 |
| Longueville | 9 | 1 | 3 | 0 | 2 | 1 | 2 | 0 |
| Orsay | 316 | 124 | 120 | 37 | 1 | 1 | 1 | 0 |
| Plougasnou & | 5 | 2 | 1 | 0 | 31 | 7 | 2 | 0 |
| Santeuil & Vexin | 39 | 8 | 2 | 5 | 89 | 55 | 17 | 0 |
| Vaucluse & Simiane | 28 | 7 | 11 | 7 | 0 | 0 | 0 | 0 |
| Bangalore, India | 7 | 4 | 1 | 0 | 1 | 1 | 0 | 0 |
| Barcelona, SP | 5 | 2 | 1 | 0 | 0 | 0 | 0 | 0 |
| Oku, Cameroon | 0 | 0 | 0 | 0 | 3 | 0 | 0 | 0 |
| Cologne, DE | 0 | 0 | 0 | 0 | 0 | 0 | 0 | 0 |
| Cambridge, UK | 0 | 0 | 0 | 0 | 1 | 0 | 0 | 0 |
| Czech Republic | 17 | 6 | 4 | 9 | 3 | 3 | 1 | 0 |
| Heidelberg, DE | 0 | 0 | 0 | 0 | 1 | 0 | 0 | 0 |
| Kazakhstan | 15 | 0 | 0 | 0 | 15 | 0 | 5 | 0 |
| Los Angeles, USA | 3 | 2 | 1 | 0 | 1 | 0 | 0 | 0 |
| Norway | 1 | 0 | 0 | 0 | 7 | 0 | 4 | 0 |
| New Zealand | 6 | 3 | 1 | 0 | 2 | 0 | 0 | 0 |
| Potsdam, DE | 0 | 0 | 0 | 0 | 0 | 0 | 0 | 0 |
| Tuscany, IT | 12 | 1 | 11 | 0 | 2 | 1 | 2 | 0 |
| Vienna, Austria | 3 | 2 | 0 | 0 | 0 | 0 | 0 | 0 |
| São Tomé | 3 | 1 | 1 | 1 | 3 | 2 | 0 | 0 |
| Shanghai, China | 7 | 3 | 4 | 0 | 9 | 0 | 3 | 0 |
| Sicily, IT | 11 | 3 | 1 | 0 | 1 | 1 | 1 | 0 |
| U Warwick, UK | 0 | 0 | 0 | 0 | 1 | 0 | 0 | 0 |
| Yerevan, Armenia | 3 | 1 | 2 | 0 | 0 | 0 | 0 | 0 |
| **total** | 556 | 188 | 173 | 83 | 204 | 84 | 48 | 0 |
| **proportion** |  | 0.34 | 0.31 | 0.15 |  | 0.41 | 0.24 | 0.00 |

**Fig 3.**
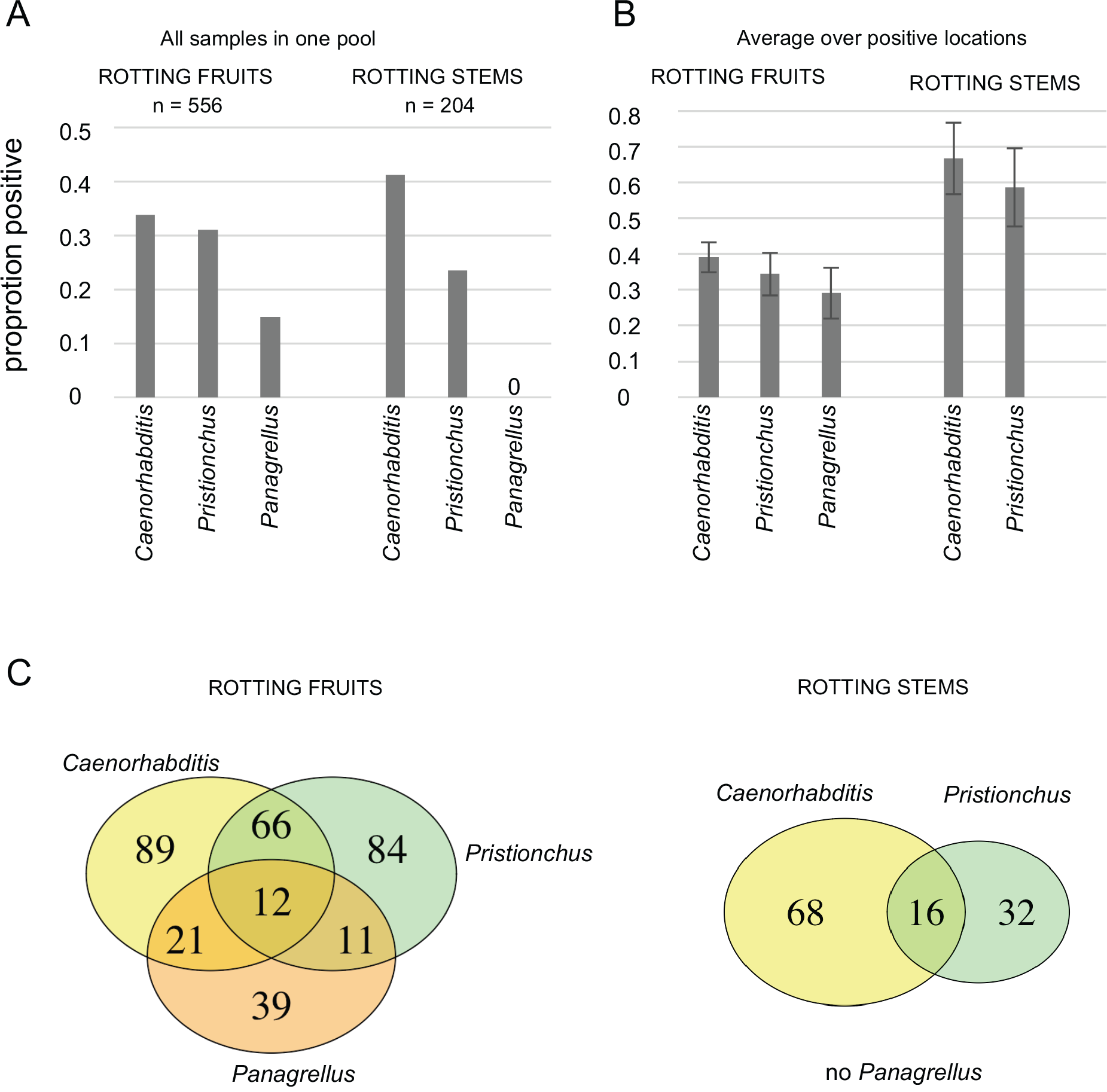
Proportions of rotting fruit and stem samples containing *Caenorhabditis, Pristionchus* or *Panagrellus*. (A) Proportions of samples positive for each genus, using all samples from rotting fruits or stems from 25 locations. (B) Proportions of samples positive for each genus, averaging for each genus over the positive locations. Bars: standard error over locations. *Caenorhabditis*, *Pristionchus* are found in both types of sample, while *Panagrellus* are found in rotting fruits only. (C) Venn diagrams for the three genera in decomposing fruits (left) and stems (right).

Among species of the *Elegans* supergroup [25], *C. elegans*, *C. briggsae*, *C. remanei* are commonly found in temperate areas. We also found *C. elegans* in samples from the Oku mountains (Cameroon) and South Yunnan (China) and *C. elegans*, *C. briggsae* and *C. tropicalis* in samples from São Tomé. Among the *Caenorhabditis* species outside the *Elegans* supergroup, we found *C. virilis* in rotting apples (Orsay, also [25]) and in droppings from a small mammal having eaten fruits (Longueville), in both cases close to Paris, France. *C. portoensis* was found in several places in Western Europe in rotting fruits: in apples in Portugal [25] and France, and in oranges in Sicily. Finally, *C. monodelphis* [55] was found in one rotting wood sample including a tree fungus, from Oslo, Norway.

### *Pristionchus* is commonly found in rotting vegetal matter

*Pristionchus* was found in 31% of rotting fruit samples (n=556) and 24% of stem samples (n=204; Table 1), thus in a comparable proportion to *Caenorhabditis*, especially considering that the numbers for *Pristionchus* are underestimates (especially in the large set of Santeuil stem samples, where *Pristionchus* was not always distinguished from other diplogastrids; S1 Table). *Pristionchus* is thus commonly found in both rotting fruit and stem samples.

In addition to rotting fruits and stems, we noted the presence of *Pristionchus* in other types of decomposed vegetal matter (S1 and S2 Tables): soil/humus and leaf litter (over 20 samples), compost, flowers, iris and hyacinth bulbs, cacti, leaves, moss, and wood. We also found *Pristionchus* in fungi (on trees or on the ground), a *Geophilus* myriapod, a dead bee, a dead *Helix aspersa* snail (S1 and S2 Tables), and droppings from a small mammal. This list is quite similar to the samples where we found *Caenorhabditis.* The only exception may be that *C. elegans* and *C. briggsae* were found in live snails, slugs and isopods (here and [21,22,27,30]) and we never found *Pristionchus* in such samples.

*Pristionchus* was found in most geographic locations (23/26 locations; while *Caenorhabditis* was found in 21/26 locations). The three locations where *Pristionchus* was not found were those sampled for *Mesorhabditis* spp. (mostly humus/rotting leaves samples), where *Caenorhabditis* was also not found. The two locations where *Pristionchus* but not *Caenorhabditis* was found were at Northern latitudes (Lofoten Islands) and on mountains with a continental climate and negative temperatures in winter (ca. 1500-2000 meters altitude near Almaty, Kazakhstan). Conversely, it would be interesting to establish whether *Pristionchus* may be less common than *Caenorhabditis* in equatorial regions (as could be suggested from samples from São Tomé and Bangalore; S1 Table).

Within a single sample, the census size of *Pristionchus* appeared comparable to *Caenorhabditis*, with populations ranging from 1-few animals to over 1,000 (over 10,000 for *Caenorhabditis*) in one sample (S1 Table; for example samples shown in Fig 1B,F). *Pristionchus* feeding stages were noted to be present alongside dauer larvae in both rotting fruits and stems (Orsay, Santeuil). *Pristionchus* was the predominant species in terms of numbers in some samples (S1 Table), while in others *Caenorhabditis*, *Oscheius*, other rhabditids, or *Panagrellus* were most abundant.

A variety of *Pristionchus* species were found, as indicated by our 18S sequencing (S2 Table), consistent with previous reports [7,8,11,56]. Among them, we found *P. pacificus*, as verified by crosses (S2 Table). Some of our 18S sequences do not match those of any *Pristionchus* species in the databases and may be new species. Conversely, some *Pristionchus* species may not be distinguished by this short fragment. Importantly, the sampled diversity covers all groups of *Pristionchus* species in [56], except the more basal ‘*Elegans*’ group. Indeed, we found 18S best hits in the *Pacificus* group to *P. pacificus*, *P. arcanus*, *P. japonicus* and *P. quartusdecimus*, in the *Maupasi* group to *P. atlanticus*, in the *Lheritieri* group to *P. uniformis* and *P. entomophagus*, and in the *Triformis* group to *P. triformis* and *P. hoplostomus*. In addition, in a rotting bulb we found a putative representative of the fresh fig clade [15] with a sequence resembling that of *Pristionchus* sp. 35.

### *Panagrellus redivivoides* is found in rotting fruits but not rotting stems

*Panagrellus* nematodes were found in 15% of rotting fruit samples (n=556; confidence interval using a binomial distribution 12-18%) and 0% of the rotting stem samples (n=204; confidence interval 0-2 %) (Table 1). Thus, in our sampled substrate types, *Panagrellus* is specifically enriched in fruits versus stems (Fisher exact test rejecting homogeneity, *p*=10^−12^). Rotting fruits contain bacteria and fungi and we observed that *Panagrellus* adults were often attracted to fungi (yeasts) on the culture plates.

Another invertebrate commonly found on these rotting fruits but not rotting stems is the fruit fly *Drosophila* (various species). We sampled live *Drosophila* in the field and on two occasions found *Panagrellus* (S2 Table and [26]). We also found *Panagrellus* in a *Drosophila willistoni* culture (S2 Table). We observed *Panagrellus* dauer larvae climbing on *Drosophila* pupae in our samples and waving, especially on the pupal appendages (Fig 2B). It is thus possible that fruitflies constitute vectors for *Panagrellus* between rotting fruit food patches.

We performed pairwise mating tests between *Panagrellus* isolates and to our surprise, found that all our fruit isolates were compatible, thus representing a single biological species. Morphologically, the vulva slit appeared almost perpendicular to the ventral side of the females (Fig 2C,D, S2B Table). This feature is characteristic of a single species of *Panagrellus*, called *P. redivivoides* [57,58]. In all other species such as *P. redivivus*, the vulva is strongly bent towards the anterior side of the animal (Fig 2E,F). Other morphological features of our isolates also matched the original and subsequent redescription of *P. redivivoides* [57,59,60], including the shape of the spicules and the gubernaculum dorsal end bearing a hook in lateral view (Fig 2G,H).

This *Panagrellus* species was collected from diverse fruit samples (apples, pears, peach, grapes, plums, cherries, tomatoes, walnuts, figs), but never from other substrates (except once in compost containing rotting fruits). Among the substrates we sampled, *P. redivivoides* thus appears specific to rotting fruits. It does not appear in every orchard or region where we have sampled, for example it was not found in Utah, while it was common in Oregon and Washington states. We recently found a new *Panagrellus* isolate (JU3343) in a compost heap of unclear composition but containing no fruits. As an exception that proves the rule, this isolate corresponds to a different *Panagrellus* species both through mating tests and morphology (S2B Table).

We amplified and sequenced 18S and 28S rDNA fragments for the *Panagrellus* isolates, which yielded the same sequence for all our rotting fruit *Panagrellus* isolates, confirming our crossing tests. The Stock and Nadler article [59] that aimed to associate DNA sequence tags to *Panagrellus* morphological species did not include live cultures of *P. redivivoides*. Two later articles with new *Panagrellus* sp. isolates [53,61] yielded closely related sequences but the authors did not attempt to identify the species morphologically or through mating tests. We found that the faster evolving 28S rDNA fragment was most similar but not identical to that of *Panagrellus* sp. MC2014 KM489128 (isolated from an aberrant specimen of the red palm weevil *Rhynchophorus ferrugineus* in [61]). The sequence of the 18S rDNA fragment is also similar but not identical to *Panagrellus* sp. MC2014 and to a newly described species *Panagrellus levitatus* [62] (see Discussion).

### Other nematodes

Other nematode genera were found in rotting fruits and stems. *Oscheius* (species of the *Dolichura* group including the common *Oscheius tipulae*) was present in all types of samples. Although it was found frequently in soil [63], we had never observed feeding stages until we started collecting rotting fruits and stems (S1 Table). Other rhabditids included *Rhabditella*, *Auanema*, *Pelodera*, etc. *Mesorhabditis* was often present in small numbers, and also found in humus and rotting leaves in forests. *Panagrolaimus* was also found in soil but often present in rotting fruits and stems, even if quite dry. *Rhabditophanes* was particularly observed in cold weathers and climates. Fruit samples also commonly contained fungi-eating nematodes such as aphelenchs.

### Co-occurrence

Many of the different nematodes co-occurred in the same sample (Table 1). Fig 3C summarizes the co-occurrence of the three genera *Caenorhabditis*, *Pristionchus* and *Panagrellus* in fruits and stems. A weak positive correlation was found in rotting fruits for the co-occurrence of *Pristionchus* and *Caenorhabditis* (correlation coefficient *r* = 0. 16 [0.08-0.24 confidence interval], *p* = 0.0001). For example, in fruits, 41.0% of the samples with *Caenorhabditis* (n=188, for a total of 556 fruits) also had *Pristionchus* and conversely, 44.5% of the samples with *Pristionchus* (n=173) contain *Caenorhabditis.* No other correlations were found significant (note however that sample sizes were smaller in stems and for *Panagrellus*).

### Seasonal pattern in the Orsay orchard

Fig 4 shows the abundance of the three genera over time in apples of the Orsay orchard. *Pristionchus* was present in rotting apples on every tested date of the year, spanning July to January. *Caenorhabditis* was not found on the two January dates (see further data in [26]) and *Panagrellus* was not found in July (twice in two different years) nor on three dates in December and January.

**Fig 4.**
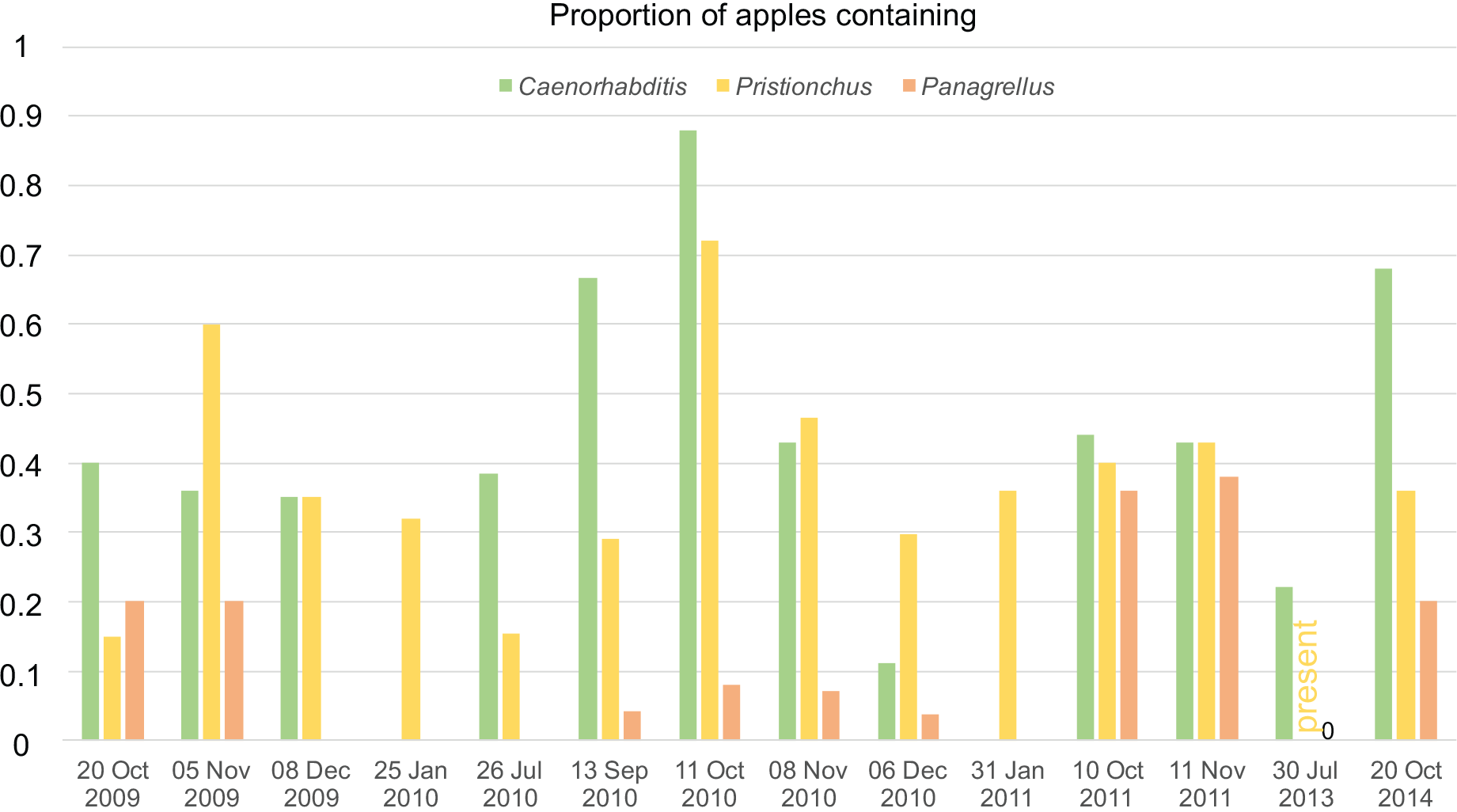
Seasonal pattern of presence of *Caenorhabditis, Pristionchus or Panagrellus* in apples of the Orsay orchard. This figure shows the proportion of positive apples for *Caenorhabditis*, *Pristionchus* or *Panagrellus* at different dates of sampling. *Pristionchus* was noted present in a few apples on the 30 July 2013 timepoint but the number of positive apples was not scored. n=20-28 apples per date.

## Discussion

### Not only fond of beetles: *Pristionchus* also lives in rotting vegetal matter

Our results demonstrate that *Pristionchus* nematodes are found in rotting vegetal matter frequently and abundantly, in both dauer and feeding stages. This includes *P. pacificus* and other “beetle-associated” *Pristionchus* species, such as *P. uniformis*, *P. entomophagus*, *P. triformis*, *P. quartusdecimus* or *P. atlanticus* [8,9,12,56,64], or closely related species (S2 Table). From these data, we conclude that these *Pristionchus* species feed in rotting vegetal matter and are not exclusively necromenic. These decomposing invertebrates are likely not their main source of food compared to the microbial blooms and other nematodes that *Pristionchus* may encounter on rotting vegetal substrates over several seasons.

From our literature review (see Introduction), *Pristionchus* have not been shown to be feeding in the wild on naturally decomposing beetle corpses. Thus, although it cannot be ruled out that *Pristionchus* may be necromenic on beetles in some instances, the relationship of *Pristionchus* species with beetles may be similar to the relationship of *Caenorhabditis* species with isopods or terrestrial molluscs. Indeed, when any of the carriers of *Caenorhabditis* are killed on a Petri dish as *Pristionchus*-bearing beetles have been, *Caenorhabditis* will exit the dauer stage and start reproducing. (Note that on live slugs and snails, other developmental stages of *C. elegans* are also found [26,27]). Evidence for a possible *C. elegans* necromenic behavior is anecdotal: non-dauer *C. elegans* were found once on a naturally dead *Helix* snail (Table 1 in [22]). Note that we found *Pristionchus* spp. on a live *Geophilus* myriapod, a dead snail and a dead honeybee (S2 Table; nematode developmental stage unknown), and some *Pristionchus* species also have been described on invertebrates other than beetles (e.g. *P. entomophagus* on a pamphilid wasp [65]). It is thus unclear for each *Pristionchus* species whether beetle species are the only specific associates. The present survey focused on rotting vegetal matter and a few other substrates and thus provides a narrow window on *Pristionchus* ecology, yet still considerably enlarging the sample diversity compared to only collecting beetles.

Each *Pristionchus* species (and possibly population) needs to be considered separately. However, our survey found a variety of *Pristionchus* species covering several clades within the genus (S2A Table), with a biogeography consistent with that in previous reports in beetles [64]. Most of our sampling work was in Europe, where *P. pacificus* is rare compared to other *Pristionchus* spp. [8,9,11,66], but we found *P. pacificus* in decomposing vegetal matter in Asia (S2 Table).

### Consequences for the life cycle of *Pristionchus*

A representation of the *P. pacificus* life cycle has recently been proposed [17], based on the beetle association and laboratory findings. In this model, adult *P. pacificus* (and possibly dauers) are attracted to the oriental beetle *Exomala orientalis*. On the beetle, eggs and dauers are produced and developmentally arrest due to a beetle chemical studied in the article. Upon beetle death, *P. pacificus* exit the dauer stage and as adults may be attracted to another beetle. Alternatively, they adopt a “free” life cycle (likely meaning that they do not go through a dauer stage) and eventually may reenter the dauer stage or be attracted to a new beetle as adults. Presumably, the free life cycle is on the dead beetle.

Several features of this proposed life cycle are surprising: i. the stage that is attracted to the beetle is the adult nematode (which has not been observed so far on wild-caught beetles); ii. once on the beetle, the adult nematode is able to produce dauer progeny. This new view of the nematode life cycle was probably derived from the fact that most behavioral studies of chemotaxis towards beetles used *Pristionchus* adults [6,17,67,68], with the exception of [69,70], which studied dauer nictation.

We propose in Fig 5 an alternative life cycle for *P. pacificus* and other *Pristionchus* species that may be found associated with various beetles. This life cycle is similar to that proposed for *C. elegans* [30,71]. In our model, rotting vegetal matter is the feeding ground for *Pristionchus* spp., whose populations expand until the food is exhausted whereupon the young larvae enter the dauer stage. Beetles may transport dauer larvae between these patchy plant food sources. In this life cycle scenario, the association of dauer larvae with the larger invertebrate is phoretic. Necromeny may be occasional, but its occurrence remains to be demonstrated.

**Fig 5.**
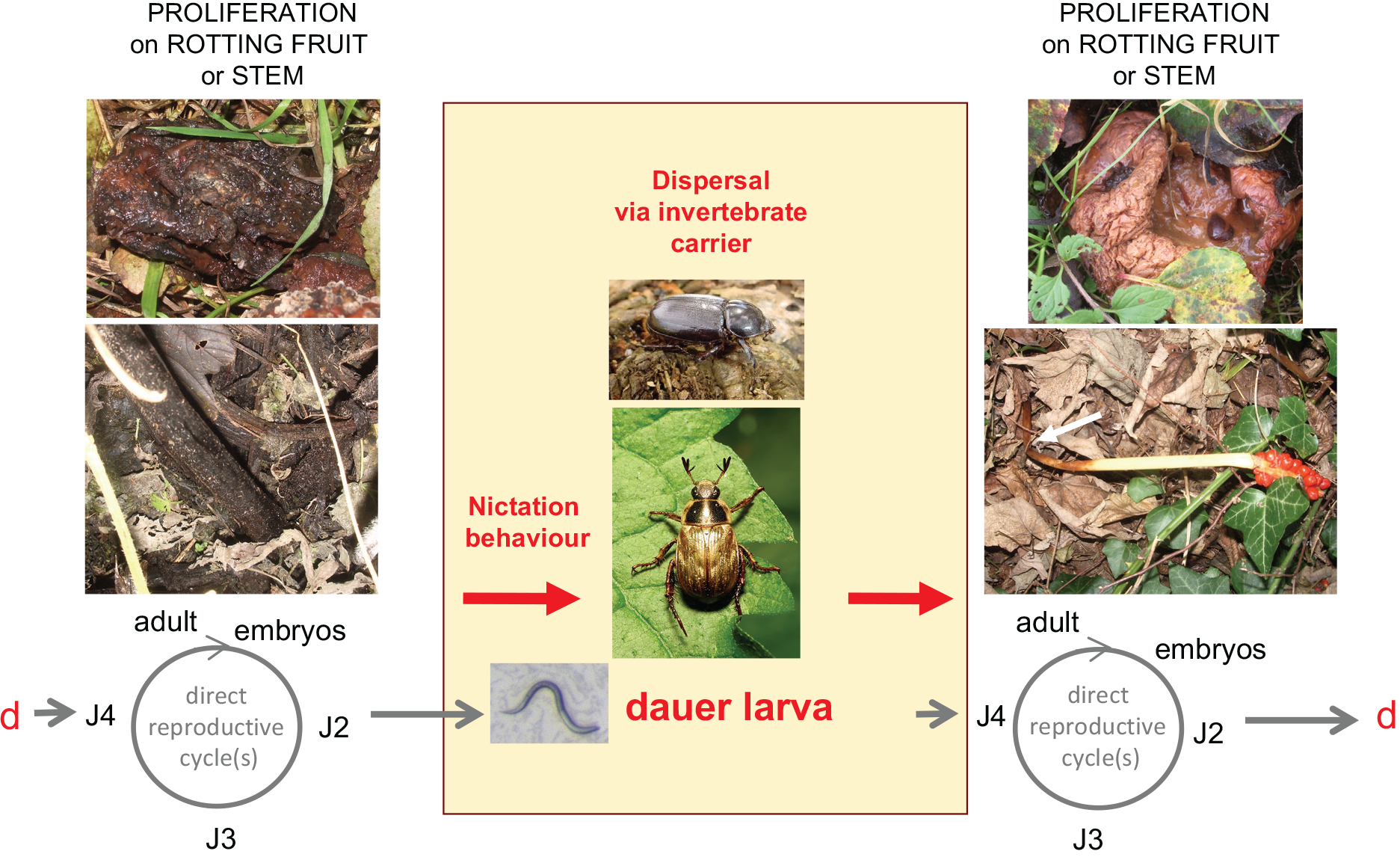
Proposed life cycle of *Pristionchus* spp. in their natural habitat. *P. pacificus* and other *Pristionchus* species proliferate in various types of rotting plant material, such as fruits. Dauer larvae are the stress-resistant, alternative third juvenile stage. (The first larval stage occurs within the embryo [97].) Dauer larvae may actively disperse to colonize new food sources. Alternatively, their nictation behaviour - standing on their tail and waving individually or in group - may allow them to attach and disperse via carriers, such as *Exomala orientalis* or *Oryctes borbonicus* for *P. pacificus*, until a new food source is encountered, where development resumes. J2-J4, juvenile stages; d, dauer larva. Modified after [71] drawn for *C. elegans*. Apples and stems: Pictures by MAF. Picture of *P. pacificus* JU1102 dauer by MAF. Bottom beetle: *Exomala orientalis*, by Katja Schulz via Wikimedia Commons CC BY-SA 2.0, https://commons.wikimedia.org/wiki/File%3AOriental_Beetle_-_Flickr_treegrow_(1).jpg. Top beetle: *Oryctes borbonicus*, by Jjargoud - Own work, CC BY-SA 3.0, https://commons.wikimedia.org/w/index.php?curid=5063614.

*P. pacificus* and other *Pristionchus* spp. may feed on bacteria, fungi and other nematodes as a food source in decomposing vegetal matter. The predatory behavior of many *Pristionchus* species towards other nematodes is rendered possible by the plasticity of development of the mouth form [15,72,73]. From our data, it is clear that many other nematodes co-occur with *Pristionchus* species in rotting vegetal matter, including but not restricted to *Caenorhabditis* and *Panagrellus* species (S1 Table). These other nematodes may compete for bacterial food, or serve as prey for *Pristionchus*. We found a weak positive correlation for the co-occurrence of *Pristionchus* and *Caenorhabditis* in rotting fruits. Rather than a specific attraction of *Pristionchus* to *Caenorhabditis-containing* substrates (or conversely), we propose that this correlation may be simply explained by the degree of decomposition, humidity or microbial fauna in the fruits, which needs to be such as to sustain these nematode species.

*C. elegans* and *P. pacificus* are the two most studied species of each genus and *C. elegans* was shown to avoid sulfolipids secreted by *Pristionchus pacificus* [74]. Due to geographical sampling biases for each genus, we have so far not detected the co-occurrence of *C. elegans* and *P. pacificus* in the same sample. It would be interesting to sample further in areas where both *C. elegans* and *P. pacificus* were found, such as Southern California, Hawaii, South Africa or La Réunion [11,75]. It is possible that the avoidance behavior of *C. elegans* may have evolved as a response to several *Pristionchus* species, and that conversely, several *Caenorhabditis* species may show avoidance to *Pristionchus pacificus* or other *Pristionchus* species.

Are the beetles that have been associated with *Pristionchus* spp. likely to visit fruits or a forest floor with rotting stems? Detailed studies would be needed to answer this important question, taking into account the seasonality of the beetle life cycle itself. *Geotrupes stercorosus* was found to be the most reliable beetle source of *Pristionchus* nematodes in Europe by [9]. Indeed, *Geotrupes* adults may feed on forest litter or fungi [76] and lay eggs in nests they provide with forest litter [77]. The *P. pacificus*-carrying *Exomala orientalis* adults emerge from soil [78] and adult females may feed on flowers [79,80]. (Whether *Pristionchus* are present on larval stages of beetles is unclear.) Indeed many of the scarab beetle species associated with *Pristionchus* were found to feed as adults on soft, high-quality diets and the micro-organisms therein [77] and many of the American tropical “dung beetle” species were found to feed on mature and rotting fruits [81]. Understanding the relationship between the rotting vegetal matter and the beetles is required to understand the biology of *Pristionchus* spp. and test our hypothesized life cycle (Fig 5).

### *Panagrellus* in rotting fruits: a specific habitat for *Panagrellus redivivoides*?

The *Panagrellus* genus was previously found on a variety of substrates. The ‘sour paste nematode’ *Panagrellus redivivus* is used in many studies because of its ease of culture in the laboratory [82–89]. This species used to be commonly found in glues used to hang wallpaper and bind books [82,90]. Other *Panagrellus* species have been found in the slime flux or cankers of trees caused by bacterial or fungal diseases, in association with bark beetles or their frass, inside pitcher plants and, as a human associate, in beer mats and spoiled cider [59,60].

Taxonomic characterization of different *Panagrellus* species can be found in [38,59,60,91], with molecular data in [59]. The vagina of all but one of the *Panagrellus* species is tilted anteriorly towards the uterus, while a vagina that extends perpendicularly to the ventral cuticle of the female is specific to *Panagrellus redivivoides* [38,59,91]. *P. redivivoides* did not yet have a molecular tag to anchor the morphological description.

Our mating tests show that all the *Panagrellus* we isolated on rotting fruits or *Drosophila* belong to a single biological species, with a morphology matching *Panagrellus redivivoides* [57,58]. Its 28S rDNA sequence is identical to that of a *Panagrellus* sp. found in decomposing pomegranate in Italy [53] and both 28S and 18S sequences are similar but not identical to that of a *Panagrellus* found in a weevil in [61]. A new species of *Panagrellus* has been very recently described from a culture of *Drosophila melanogaster* and called *P. levitatus* [62]. This species presents similar but not identical 18S rDNA sequences to those we found (S2 Table) and has a weakly tilted vulva. The only morphological character of this isolate that is supposed to differ from *P. redivivoides* is the presence of circumcloacal papillae in *P. levitatus,* with a reference to [91] for their “absence” in *P. redivivoides* (to our knowledge only an absence of information). Further work would be required to make sure that there are two different species and because of differences in rDNA sequence with *P. levitatus*, abundance on several continents and priority, we identify the biological species we found in rotting fruits as *Panagrellus redivivoides*.

Concerning phoretic associations, *P. redivivoides* may be associated with *Drosophila* fruitflies, in contrast to other *Panagrellus* sp. that appear associated with bark beetles. The fruifly association relies on several reports. Early on, using *Drosophila* traps made of potato puree, Aubertot [92] found “*Panagrellus silusiae*”, now synonymized with *P. redivivus* [90,93] (yet the species determination of Aubertot is unclear, especially since the species *P. redivivoides* was not described yet). *P. redivivoides* was found several times in laboratory cultures that were visited by fruitflies, including in the original description of the species by Goodey [57], which reports the arrival of the species in a banana maize-meal cider used to trap *Drosophila* flies. Lees [82] set up experiments to test the association of what he called “*P. silusiae*” (again, the species identification is unclear) and *Drosophila funebris* and showed that the latter could transport the former to new Petri dishes. “*Panagrellus zymosiphilus*”, now synonymized with *P. redivivoides* [91] was also found twice in *Drosophila* cultures, the second time on *Drosophila obscuroides* recently isolated from nature [94,95]. “*P. zymosiphilus*” was also found in grapes, presumably brought by a *Drosophila* [96]. We added one data point to this association with *Drosophila* cultures with JU385, isolated in a *Drosophila willistoni* culture (S2B Table). In addition, *Drosophila* larvae are very often found in the rotting fruits we sampled and we could isolate *Panagrellus* sp. on *Drosophila* caught outside in two locations, Orsay and Le Blanc (reported in [26]), the latter yielding JU1055 (S2B Table).

We suggest the possibility that one and the same species, *P. redivivoides*, is the most commonly associated with rotting fruits and *Drosophila* larvae, a distinguishing ecological feature compared to most *Panagrellus* species. Its life cycle may be similar to that depicted for *C. elegans* [71] or *Pristionchus* (Fig 5), but with *Drosophila* fruitflies carrying the dauer larvae from fruit to fruit.

In summary, based on our sampling and the observed distribution of feeding and dauer stages, we propose a life cycle for *Pristionchus* nematodes and *Panagrellus redivivoides* that is similar to that of many *Caenorhabditis* species, including *C. elegans*, whereby they feed on the microbial blooms on decomposing vegetal matter and are transported between these food patches by larger invertebrates, which may be beetles for *Pristionchus* spp., fruitflies for *Panagrellus redivivoides* and isopods and terrestrial molluscs for *Caenorhabditis* spp.

## Acknowledgements

We thank members of our labs and all other sample collectors listed in Tables S1 and S2, especially Jim Thomas. We thank Irini Topalidou for assistance with PCR and sequencing. We thank C. Braendle, S. Chalasani, B. Schlager and H. Teotonio for reading the manuscript.

## Supplementary information captions

**Table. Surveys of some nematode genera in various sample types from different worldwide locations.** Each sheet corresponds to a sampling location and is designated by a letter: A-G, locations in France, H-Z locations outside France, in alphabetical order. Presence of a nematode genus is indicated by ‘x’. When the relative abundance of different genera has been scored, 1 indicates the most abundant, followed by 2, 3, etc. *Caenorhabditis* (green column), *Pristionchus* (yellow), *Panagrellus* (orange) are indicated first. The other groups are indicated roughly in order of ease of extraction from the sample and identification. On the right, in columns ‘*Caenorhabditis* abundance log index’ and ‘*Pristionchus* abundance log index’, is indicated the abundance log index (see Methods): here 5 is highest and 1 lowest. ‘f’ indicates feeding stages, ‘d’ dauer larvae, in order of abundance, with the caveat that dauer larvae are more difficult to identify than feeding stages. Note that the care and timing with which the different nematodes were identified and monitored for number and stages differ among the different sampling dates and locations. Some data for *Caenorhabditis* are from [26,49]. *Pristionchus* are best distinguished from other diplogastrids in the adult stage; thus, a ‘*’ in the ‘other diplogastrid’ column indicates that these diplogastrids could have been *Pristionchus*. The *Caenorhabditis* species are abbreviated ‘Cel’ for *C. elegans*, ‘Cbr’ for *C. briggsae*, ‘Cre’ for *C. remanei*, ‘Ctr’ for *C. tropicalis*, ‘Cvi’ for *C. virilis*, ‘Cpo’ for *C. portoensis*, ‘Cmo’ for *C. monodelphis*. Latitude and longitude coordinates are indicated with the number of digits corresponding to the precision.

**Table. Wild isolates from various sample types.** Sheet A: *Pristionchus*. The table indicates the origin of the strains of *Pristionchus* (with JU or BRC standard strain names), and other samples that were noted to contain *Pristionchus*. The sample ID is a temporary ID for a given collection date. Other non-*Pristionchus* frozen strains from the same samples are indicated on the right. nd: not determined. Sheet B: *Panagrellus*. The table indicates the origin of *Panagrellus* strains. Sheet C: Other species of nematodes collected from rotten fruit and characterized by 18S rDNA sequencing.

